# *ABCA7* rs3752231 variant effects on glial responses to amyloid-β and plaque maturation in Alzheimer’s disease

**DOI:** 10.64898/2026.06.18.733103

**Authors:** Marianna Papageorgopoulou, Emily Adair, Nurun N Fancy, Baptiste Avot, Samuel L Boulger, Mara S Wülfing, Paul M Matthews

## Abstract

Variants in *ABCA7* are among the most consistently replicated genetic risk factors for late-onset Alzheimer’s disease (AD), yet the cellular mechanisms remain poorly defined. Here, we characterise the impact of the common *ABCA7* rs3752231 risk variant on amyloid-β (Aβ) pathology and glial responses in human *post-mortem* brain, combining quantitative neuropathology of 4G8-immunostained mid-temporal gyrus from 99 donors (Braak 0-VI) with glial-enriched single-nucleus RNA sequencing from 54 of them. *ABCA7* rs3752231 carriers exhibited an increased Aβ burden and larger plaques with late AD explained by a selective expansion of diffuse plaques and relative reduction in compact plaques, consistent with impaired microglial-mediated plaque maturation. Transcriptional responses to increasing Aβ burden were largely genotype-specific: non-carriers showed canonical disease-associated microglial activation, including upregulation of complement, phagocytic, and inflammatory pathways, alongside astrocyte responses consistent with preserved synaptic support, while carriers exhibited a distinguishable activation state. Exploratory ligand-receptor analysis identified carrier-specific intercellular signals suggesting non-cell autonomous suppression of microglial phagocytosis. Together, these findings position *ABCA7* rs3752231 as a regulator of glial responses to AD pathology, linking a common coding variant to impaired microglial plaque containment and maladaptive astrocyte responses and nominate microglial *TREM2* activation and *CD33* inhibition and astrocytic EAAT2 induction as candidate therapeutic strategies.

## Introduction

Alzheimer’s disease (AD) is the most common cause of dementia, defined neuropathologically by extracellular amyloid-beta (Aβ) and intraneuronal phosphorylated tau accumulation accompanied by neuroinflammation and progressive neurodegeneration ^1,2,3^. The vast majority of cases are late-onset and arise from a polygenic and multifactorial genetic architecture; genome-wide association studies (GWAS) have identified more than 70 risk loci for late-onset AD expressed in glio-vascular vascular cells and predominantly by microglia ^4,5^. These implicate particularly microglial innate immune and mechanistically associated lipid metabolism and transport as central contributors to disease pathogenesis^4,5^.

*ABCA7* arguably has the greatest allelic heterogeneity of any AD risk locus other than *APOE*; with both rare and common *ABCA7* variants implicated^6^. The rs3752231 variant is a common (alternate allele frequency ∼0.33), missense mutation that confers a moderate AD risk (odds ratio = 1.09)^6^. The gene encodes a transmembrane ATP-dependent lipid transporter, expressed across brain cell types, with enrichment in microglia^7^. The rs3752231 variant acts through haploinsufficiency^8^. In mouse models, *Abca7* haplodeficiency impairs inflammatory responses to lipopolysaccharide stimulation *in vivo* and disrupts *CD14* expression^9^, while *Abca7* deficiency in mouse AD models exacerbates Aβ burden and impairs microglial responses to pathology^10^. *In vitro* studies using human macrophage lines demonstrate that *ABCA7* determines phagocytic capacity^11^ and human CSF data from carriers of *ABCA7* mutations provide evidence for reduced inflammatory biomarkers and altered APP processing^12^. These observations raise the question of whether *ABCA7* variation influences not only the quantity but also the morphology of Aβ deposits in the human brain, a distinction with direct implications for microglial functional competence, Aβ clearance and compaction. A recent snRNA-seq study of different loss-of-function *ABCA7* variants in cortical tissue identified transcriptomic changes across multiple cell types, with enrichment in excitatory neurons^13^. However, whether neuronal dysfunction relevant to the genesis of AD can be attributed primarily to the cell autonomous loss of function of ABCA7 and how disease risk is conferred by missense variant alleles in microglia (or other glia) are unknown. Epigenetic profiling suggests a primary microglial influence as common genetic risk variants are disproportionately enriched in microglial enhancers^14^. Our work here will address how the common rs3752231 missense variant influences microglial responses in AD, exploring biological questions central to pathogenesis and distinct from those addressed by the prior loss-of-function studies focusing on its functional consequences for excitatory neurons^13^.

Aβ plaques exhibit marked morphological heterogeneity with distinct pathological consequences^15^. Microglia actively remodel plaques, forming a barrier that limits Aβ-associated neurotoxicity^16^. Impaired microglial activation, such as is associated with *TREM2* haplodeficiency or loss of function variants, leads to more diffuse plaques and increased neuritic damage^17,18^. Plaque morphology therefore reflects microglial functional competence rather than simply Aβ burden^19,20^. Whether common *ABCA7* variation is associated with impairment of plaque formation in human tissue and AD has not been investigated.

We have performed an integrated analysis of *post-mortem* human mid-temporal gyrus (MTG) tissue from 102 individuals spanning Braak stages 0-VI, stratified by *ABCA7* rs3752231 carrier status. Combining deep learning-assisted quantitative plaque phenotyping with single-nucleus RNA sequencing following glial enrichment, we show that *ABCA7* risk is associated with impairment of microglial and astrocyte responses to Aβ pathology. These findings implicate impaired glial engagement with amyloid pathology as a central mechanism linking common *ABCA7* variation to disease risk and identify candidate targets for genotype-stratified therapeutic intervention.

## Results

### *ABCA7* rs3752231 carriers show increased Aβ plaque burden at late disease stages

Total plaque coverage (proportion of ROI area occupied by plaques; our measure of Aβ burden) and mean plaque size were quantified across Braak stages in 99 individuals (63 carriers, 36 non-carriers; Fig. 1a). Groups were well-matched across key demographic and clinical variables. At late Braak stages (V-VI), carriers showed 1.9-fold greater total plaque coverage than non-carriers (median, 0.114 vs 0.061; Benjamini-Hochberg (BH)-corrected p = 9.09 × 10⁻⁴; Fig. 2a) and 1.6-fold larger mean plaque size (median, 1,211 vs 767 μm²; BH-corrected p = 0.010; Fig. 2b). No significant differences were observed at early (Braak 0-II) stages. The mid Braak group was underrepresented in non-carriers (n=2), precluding meaningful between-group comparison at this disease stage. The absence of differences at earlier stages suggests a progressive divergence in Aβ containment, processing and sequestration with disease progression rather than an early acceleration of deposition. Plotting Aβ coverage against individual Braak stage further illustrated this divergence, with carriers showing apparently steeper Braak-associated increases in both total and diffuse Aβ coverage than non-carriers.

**Fig. 1.**
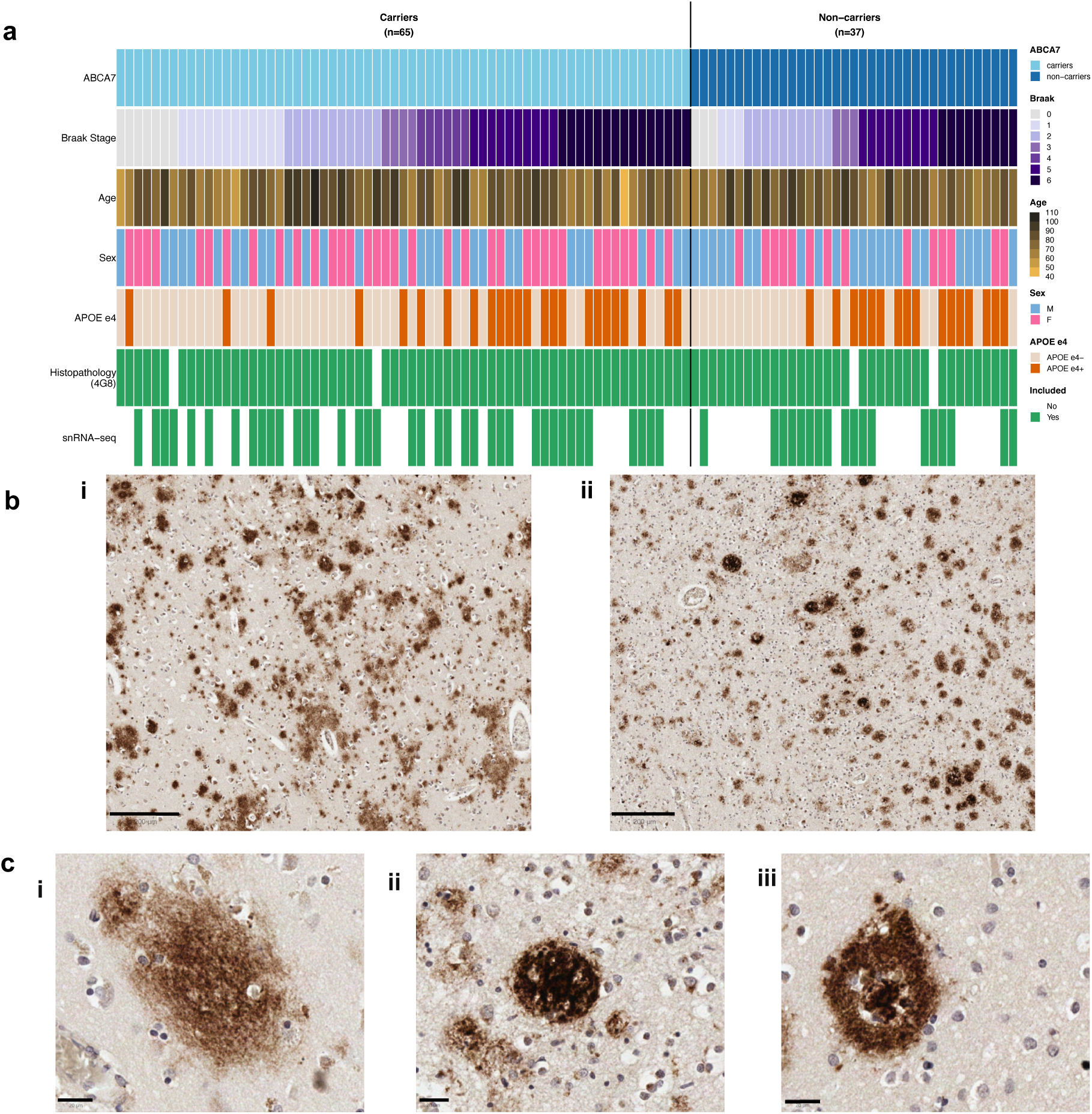
Cohort assembly and Aβ plaque characterisation in the middle temporal gyrus. Formalin-fixed paraffin-embedded (FFPE) MTG tissue was obtained from 99 donors spanning the full Braak^3^ neurofibrillary tangle staging spectrum (Braak 0-VI). **a** Cohort overview displaying demographic and neuropathological characteristics of all donors. Each column represents one individual, ordered by ABCA7 genotype (carriers, n=65, non-carriers n=37; 99 underwent IHC and 54 snRNA-seq) and Braak stage within each group. Carriers were heterozygous for the rs3752231 variant, non-carriers carry the common variant. Rows indicate Braak stage (light to dark purple, 0-6), age (light yellow to black, 46-103 years), sex (blue, male; pink, female), APOE ε4 status (orange, ε4+; cream, ε4−), histopathological assessment (4G8 immunohistochemistry; green, included) and snRNA-seq profiling (green, included). Groups were comparable for age, sex, Braak stage distribution, APOE ε4 status, and post-mortem interval (Mann-Whitney U test for continuous variables; Fisher’s exact test for categorical variables). **b** Aβ immunohistochemistry from an ABCA7 carrier (**i)** and a non-carrier **(ii)**, illustrating the greater size and density of diffuse plaques in carriers in a representative MTG cortical field. Scale bars, 200 µm. **c** Representative high-magnification images illustrating the three plaque morphological subtypes: diffuse **(i)**, characterised by poorly defined amorphous Aβ deposits; compact (ii), with a well-defined rounded boundary and uniform dense Aβ; and dense-core **(iii)**, comprising a compact Aβ core surrounded by a diffuse peripheral halo. Scale bars, 20 µm.

**Fig. 2.**
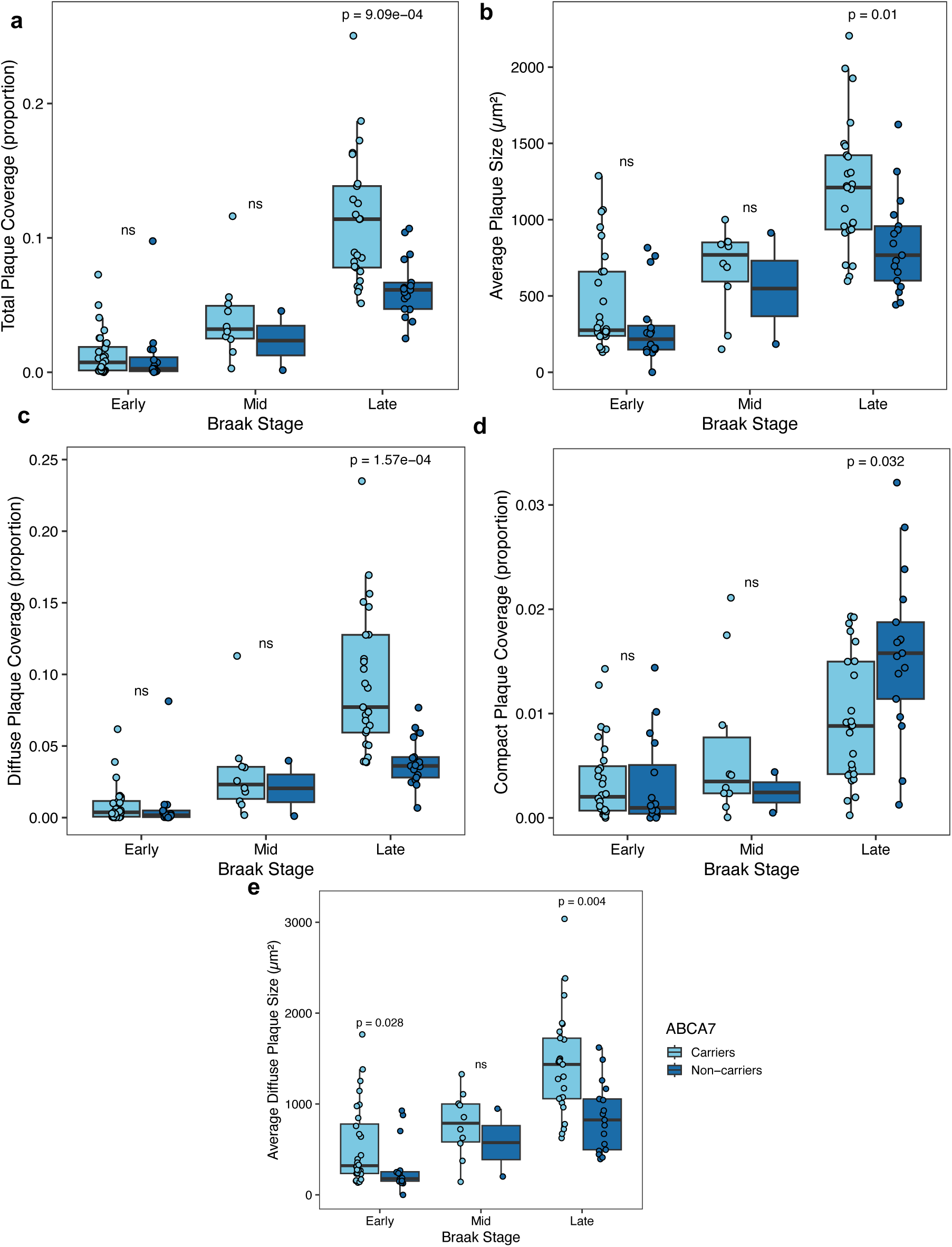
ABCA7 rs3752231 carriers display greater Aβ plaque burden than non-carriers at late Braak stages. Quantitative histopathological analysis of Aβ plaque burden across early (Braak 0-II; carriers=28, non-carriers=16), mid (Braak III-IV; carriers=10, non-carriers=2), and late (Braak V-VI; carriers=25, non-carriers=17) disease stages in ABCA7 rs3752231 carriers and non-carriers. Light blue, carriers; dark blue, non-carriers. Box plots show median with interquartile range; individual data points represent single donors. **a** Total plaque coverage expressed as a proportion of the region of interest (ROI) area. **b** Average plaque size (μm²) across all morphological subtypes. **c** Diffuse plaque coverage as a proportion of ROI area. **d** Compact plaque coverage as a proportion of ROI area. **e** Average diffuse plaque size (μm²). Between-group comparisons at each Braak stage were performed using Wilcoxon rank-sum tests with BH correction applied across all 15 comparisons (5 measures × 3 Braak Stages); corrected p-values are shown above each comparison; ns, not significant.

### *ABCA7* rs3752231 carriers show increased diffuse and reduced compact Aβ plaque burden

Plaques were classified into diffuse, compact, and dense-core subtypes by a machine learning model pre-trained on an independent cohort of 4G8 immunostained tissue (Fig. 1b, c). Classifications were manually confirmed. At late Braak stages, diffuse plaque coverage was 2.1-fold greater in carriers than non-carriers (median proportion of the region of interest (ROI) area, 0.077 vs 0.036; BH-corrected p = 1.57 × 10⁻⁴; Fig. 2c) with 1.7-fold larger diffuse plaques (median, 1,436 vs 824 μm²; BH-corrected p = 0.004; Fig. 2e). Diffuse plaques were also larger in carriers at early Braak stages (1.8-fold; median, 321 vs 180 μm²; BH-corrected p = 0.028; Fig. 2e). Compact plaque coverage was 1.8-fold greater in non-carriers at late stages (median proportion of the region of interest (ROI) area, 0.016 vs 0.009; BH-corrected p = 0.032; Fig. 2d). The mid Braak group was underrepresented in non-carriers (n=2), precluding meaningful between-group comparison at this disease stage. No differences in dense-core plaque coverage were observed at any Braak stage. Together, these results indicate that increased Aβ burden in carriers is associated with shifts in plaque subtype composition, consistent with altered plaque maturation dynamics.

### Microglial transcriptional responses to Aβ differ by *ABCA7* genotype

Our analysis here has concentrated on microglia and astrocytes because of the rs3752231-associated plaque neuropathology and the primary role of microglia in plaque formation^19,20^.Pseudobulk differential gene expression analysis was performed on 43,262 microglial nuclei enriched by negative fluorescence sorting (see Methods) from 54 donor brains (35 carriers, 19 non-carriers; Fig. 1a). Gene expression was modelled as a function of total Aβ plaque coverage (proportion of ROI covered by plaques) separately for microglial nuclei of rs3752231 carriers and non-carriers. Of 872 significant (FDR-adjusted p ≤ 0.05 and |log₂FC| ≥ 0.10) DEGs identified, only 85 (9.7%) were shared between genotypes, of which 62 showed concordant and 23 discordant directions of effect (Fig. 3a, b). The 62 concordantly differentially regulated DEGs included transcripts encoding proteins involved in lipid biosynthesis (*LTC4S, ACSL1*) and those involved in cytoskeleton-remodelling (*CADM1*, *CARMIL1*, *GSN*). *ABCA7* mRNA expression did not differ significantly between carriers and non-carriers in microglia (log₂FC = −0.16, p = 0.856) or astrocytes (log₂FC = +0.36, p = 0.664), consistent with the missense nature of rs3752231, which is predicted to alter protein stability and function rather than transcript abundance ^13^. The observed differences in transcriptional signatures in microglia for the two genotypes likely reflect downstream functional consequences of the variant protein.

**Fig. 3.**
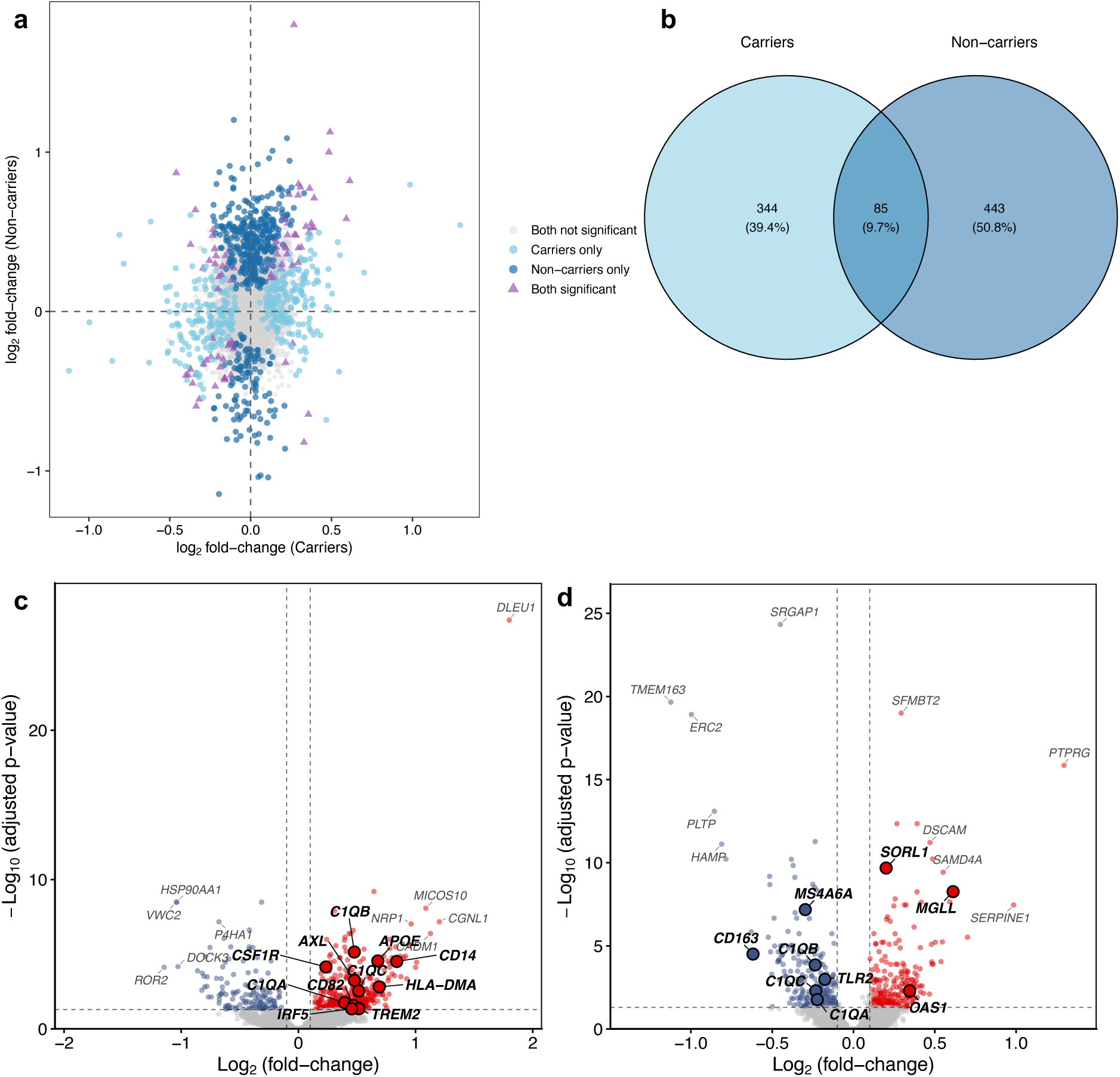
Microglial transcriptional responses to Aβ burden differ by ABCA7 genotype. Pseudobulk analysis of differentially expressed genes (DEGs) in microglia associated with greater total Aβ plaque coverage, identified separately in ABCA7 rs3752231 carriers (n=35) and non-carriers (n=19). DEGs were defined as adjusted p ≤ 0.05, |log₂FC| ≥ 0.10. **a** Scatter plot comparing log₂ fold-change estimates from pseudobulk models in carriers (x-axis) and non-carriers (y-axis) for all tested genes highlighting the largely distinct transcriptional responses for carriers and non-carriers. Genes significant in non-carriers only are shown in dark blue; genes significant in carriers only in light blue; genes significant in both in purple triangles; non-significant genes in grey. Concordantly upregulated genes occupy the upper-right quadrant; concordantly downregulated genes the lower-left; discordant genes occupy the upper-left and lower-right quadrants. **b** Venn diagram illustrating the simple overlap of significant microglia DEGs between carriers and non-carriers. Of 872 DEGs, 344 (39.4%) were exclusive to carriers, 443 (50.8%) exclusive to non-carriers, and 85 (9.7%) were shared. **c** Volcano plot of microglia DEGs in non-carriers with increasing Aβ burden with the top 10 genes by combined score (-log10(padj) × |log2FC|) and selected genes labelled. Red points, upregulated transcripts; blue points, downregulated transcripts; bold points discussed in text; grey points, non-significant. Dashed lines indicate significance thresholds. **d** Volcano plot of microglia DEGs in carriers with increasing Aβ burden, labelling as in **(c)**. Y-axis limits differ between panels **(c)** and **(d)** to reflect different ranges of adjusted p-values between genotype groups; axis ranges are indicated on each panel.

Non-carriers exhibited a predominantly upregulated transcriptional response (390 transcripts upregulated, 138 downregulated), including increased expression of complement (*C1QA*, *C1QB*, and *C1QC*), phagocytic receptor (*AXL*, *CSF1R*), and disease-associated microglial (DAM) marker^21^ (*TREM2*, *HLA-DMA*, *IRF5*, *APOE*; Fig. 3c) genes. In carriers, the transcriptional response was more balanced (217 up, 212 down), with upregulation of genes including *SORL1*, *MGLL*, and *OAS1* (Fig. 3d). Transcripts encoding complement components *C1QA*, *C1QB*, and *C1QC* were downregulated in carriers. A post-hoc power analysis (Parametric bootstrap, 1,000 iterations; 80% power threshold; effect sizes derived from observed limma t-statistics) suggested that the n=19 non-carrier cohort would have a high likelihood of detecting the magnitudes of effect expected for DAM markers and that the absence of equivalent enrichment in carriers therefore must reflect substantially smaller effect sizes. We conclude from this *ABCA7* rs3752231 carrier status associates with a distinct microglial transcriptional response to Aβ burden that is characterised by the absence of complement and immune-related pathways and distinct lipid metabolic and cytoskeletal process pathway signatures.

### *ABCA7* genotype modulates microglial inflammatory states and pathways associated with responses to *A*β

We extended our analysis of genotype-specific microglial signatures using an over-representation analysis of microglial DEGs (DEGs; adjusted p ≤ 0.05, |log₂FC| ≥ 0.25) for curated microglial gene sets^22^ associated with distinguishable homeostatic, disease-associated and antigen presenting cell states (Fig. 4a). In non-carriers, upregulated DEGs were enriched for HLA antigen presentation genes (adjusted p < 0.001). Antigen processing and MHC class II binding pathways were concordantly upregulated in a paired pathway analysis (Fig. 4b). In carriers, downregulated DEGs were enriched for cytokine-response (CRM2; adjusted p < 0.001) and interferon-response (IRM; adjusted p < 0.01) gene sets, consistent with an impaired or alternative activation in response to Aβ.

**Fig. 4.**
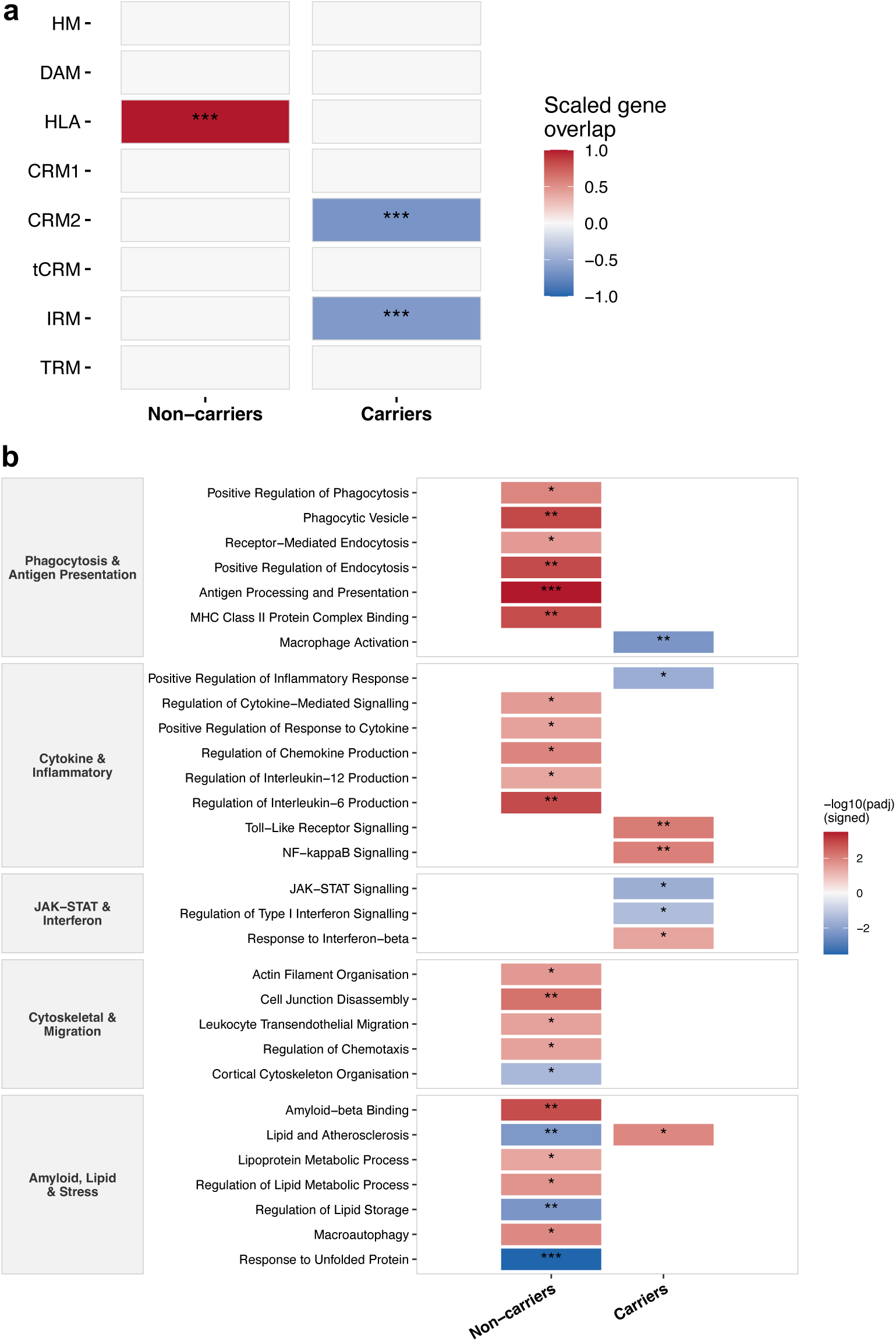
ABCA7 rs3752231 carriers and non-carriers show differences in microglial state and pathway enrichment. Enrichment analyses of gene set (a) or pathway enrichments of significantly differentially expressed microglial genes (DEGs; adjusted p ≤ 0.05, |log₂FC| ≥ 0.25) with greater Aβ burden for ABCA7 rs3752231 carriers and non-carriers. Red tiles indicate upregulated enrichment; blue tiles indicate downregulated enrichment; white tiles indicate non-significant results. Asterisks denote significance thresholds: * adjusted p < 0.05, ** adjusted p < 0.01, *** adjusted p < 0.001. **a** Over-representation analysis (ClusterProfiler) of microglial DEGs (adjusted p ≤ 0.05, |log₂FC| ≥ 0.25) for curated microglial state gene sets^22^. Tile colour represents scaled gene overlap signed by direction of regulation, ranging from −1.0 (fully downregulated overlap) to +1.0 (fully upregulated overlap). All tested states are shown. **b** Pathway enrichment analysis of microglial DEGs across five functional categories based on the Gene Ontology Biological Process, Cellular Component, Molecular Function, and KEGG databases (EnrichR). Tile colour represents −log₁₀(FDR) signed by direction of enrichment. Pathways are grouped by biological themes.

Pathway enrichment analyses (DEGs; adjusted p ≤ 0.05, |log₂FC| ≥ 0.25) revealed different patterns of enrichment for immune-related pathways between genotypes (Fig. 4b). In non-carriers, upregulated pathways included those for lipoprotein metabolic processes (*APOE*, *CTSD*), positive regulation of phagocytosis (*TREM2*, *AXL*, *C3*), receptor-mediated endocytosis, regulation of chemokine production (*APP*, *CSF1R*, *TREM2*, *IL18*), and interleukin-12 production (*CLEC7A*, *IRF5*). In carriers, canonical innate immune programmes including macrophage activation (*FCGR3A, CD93, TLR4*), positive regulation of inflammatory response (*SERPINE1, WNT5A, S100A9*), JAK-STAT signalling (*IL4R, JAK3, PTPN2*), and type I interferon signalling (*OAS1, WNT5A, PTPN2*) were downregulated. Instead, an alternative inflammatory state characterised by enrichment for toll-like receptor (*MAPK10, JUN, MAP3K8*) and NF-κB signalling (*IRAK2, BCL3, CARD11*), response to interferon-β (*PLSCR1, OAS1, XAF1*) and lipid and atherosclerosis (*MAPK10, JUN, SOD2*) pathway gene expression was upregulated (Fig. 4b). Together, these findings demonstrate that *ABCA7* rs3752231 carriers fail to adopt a canonical phagocytic microglial state in response to Aβ burden but instead express a distinct microglial transcriptional state associated with a TLR-MAPK/NF-κB-driven inflammatory signature.

### Astrocyte transcriptional responses to increasing Aβ load differ by *ABCA7* genotype

Astrocytes contribute with microglia to Aβ clearance and express *ABCA7*^23,24^. To assess how astrocyte responses to Aβ pathology differ by rs3752231 genotype, pseudobulk differential expression analysis was performed on transcriptomes from 109,122 astrocytic nuclei enriched by negative fluorescence sorting (see Methods) from 54 donors (35 carriers, 19 non-carriers; Fig. 1a), modelling gene expression as a function of total Aβ plaque coverage separately in each genotype group. Of 1,949 DEGs identified (FDR-adjusted p ≤ 0.05, |log₂FC| ≥ 0.10), 362 (18.6%) were shared between genotypes, with 332 showing concordant directions of effect (Fig. 5a, b). Both genotypes showed upregulation of previously reported reactive astrocyte markers^25^ including *GFAP*, *AQP4*, *GJA1* and the complement gene *C3* (Fig. 5c, d).

**Fig. 5.**
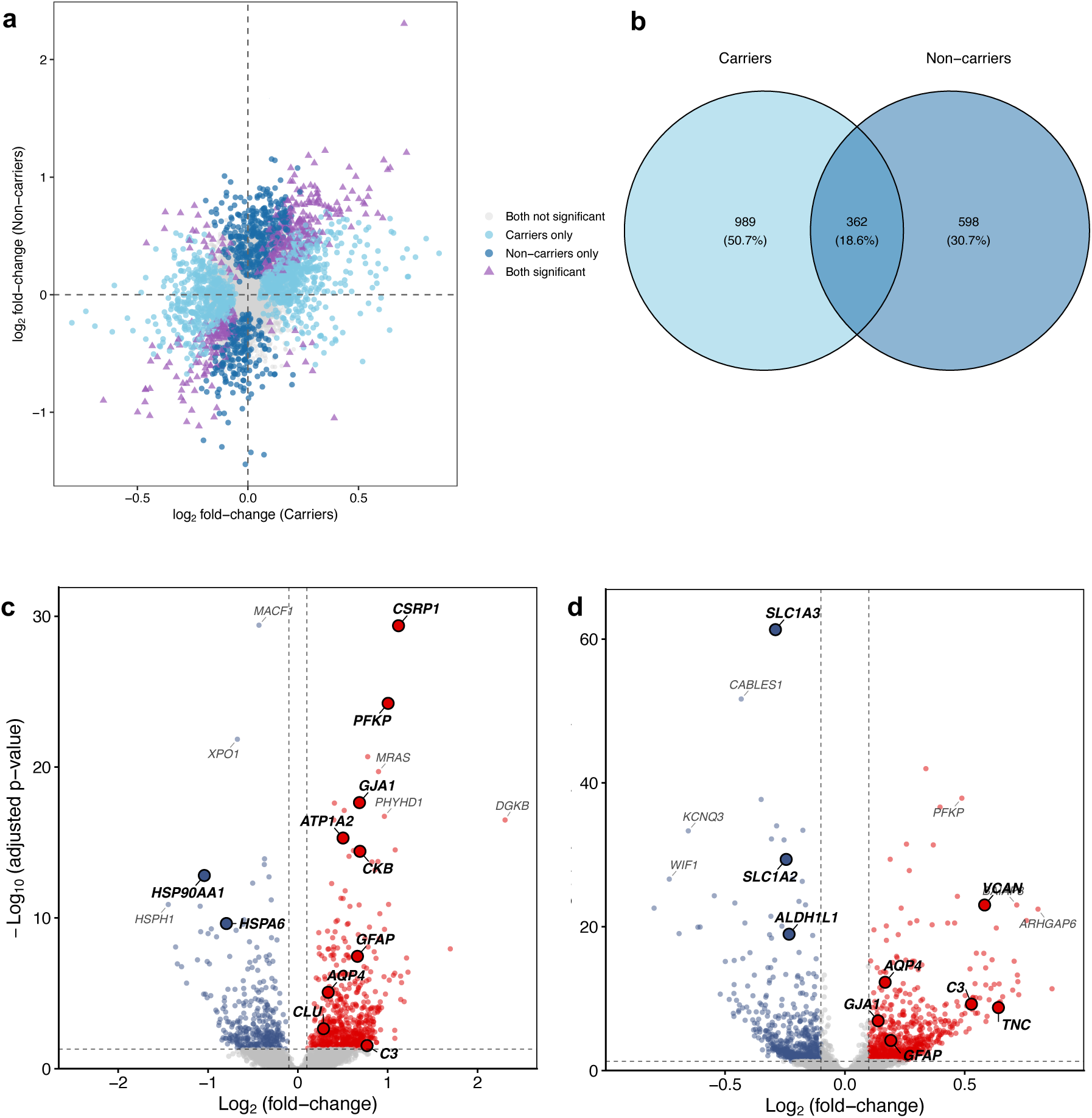
Astrocyte transcriptional responses to Aβ burden are largely distinct between ABCA7 rs3752231 carriers and non-carriers. Pseudobulk differential gene expression analysis modelling astrocyte transcriptomes against total Aβ plaque coverage, performed separately in ABCA7 rs3752231 carriers (n=35) and non-carriers (n=19). DEGs were defined as adjusted p ≤ 0.05, |log₂FC| ≥ 0.10. a Scatter plot comparing log₂ fold-change estimates from pseudobulk models in carriers (x-axis) and non-carriers (y-axis) for all tested genes. Genes significant in non-carriers only are shown in dark blue; genes significant in carriers only in light blue; genes significant in both in purple triangles; non-significant genes in grey. Concordantly upregulated genes occupy the upper-right quadrant; concordantly downregulated genes the lower-left; discordant genes occupy the upper-left and lower-right quadrants. b Venn diagram illustrating the overlap of significant astrocyte DEGs between carriers and non-carriers. Of 1,949 DEGs, 989 (50.7%) were exclusive to carriers, 598 (30.7%) exclusive to non-carriers, and 362 (18.6%) were shared. c Volcano plot of astrocyte DEGs in non-carriers with increasing Aβ burden; Top 6 genes by combined score (-log10(padj) × |log2FC|) and selected genes are labelled. Red points, upregulated DEGs; blue points, downregulated DEGs; bold points discussed in text; grey points, non-significant. Dashed lines indicate significance thresholds. d Volcano plot of astrocyte DEGs in carriers with increasing Aβ burden, labelling as in (c). Y-axis limits differ between panels (c) and (d) to reflect the different ranges of adjusted p-values between genotype groups; axis ranges are indicated on each panel.

Genotype-specific responses showed substantial differences beyond this shared small, shared gene set. In non-carriers, uniquely upregulated genes included the cytoskeletal regulator *CSRP1*, the glycolytic enzyme *PFKP*, and the energy metabolism gene *CKB*, while heat shock proteins *HSP90AA1* (log₂FC = −1.041), and *HSPH1* were downregulated (Fig. 5c). In carriers, the glutamate transporters *SLC1A3* (EAAT1) and *SLC1A2* (EAAT2), which together account for most synaptic glutamate clearance in the adult brain and reduced expression of which has been linked to excitotoxic neuronal death^26,27^, were among the most significantly downregulated genes. Transcriptional expression of the astrocyte metabolic marker *ALDH1L1* gene also was downregulated, while extracellular matrix genes *TNC*, an activator of the innate immune response and possible adaptive response to impaired microglial function^28^, and *VCAN*, also a potentiator of innate immune signalling^29^, were upregulated (Fig. 5d). A post-hoc power analysis (Parametric bootstrap, 1,000 iterations; 80% power threshold; effect sizes derived from observed limma t-statistics) confirmed that the carrier-specific downregulation of *SLC1A3* and *SLC1A2* and upregulation of *TNC* and *VCAN* were detectable with high confidence at the current carrier sample size (n=35), while the absence of the non-carrier-specific *CKB* signal in carriers reflects genuine biological attenuation rather than insufficient detection power. These findings indicate that *ABCA7* rs3752231 carrier status is associated with reduced astrocyte glutamate transporter expression alongside extracellular matrix remodelling in relation to Aβ burden, implicating impaired excitatory neurotransmitter clearance as a potential mechanism of carrier-specific neuronal vulnerability.

### Different astrocyte state enrichment and pathway-level transcriptional signatures are associated with the two *ABCA7* genotypes

Over-representation analysis using curated astrocyte state gene sets^25^ restricted to robustly perturbed transcripts (DEGs; adjusted p ≤ 0.05, |log₂FC| ≥ 0.25), revealed both shared and genotype-specific patterns of state enrichment (Fig. 6a). Reactive states astR0, astR1 and astR2 were enriched among upregulated DEGs associated with both genotypes, defining a common reactive astrocyte response to Aβ burden. The astTinf state (a growth-factor-responsive state enriched in TGF-β and trophic signalling genes) also was strongly enriched among downregulated DEGs in both genotypes (adjusted p < 0.001). Genotype-specific patterns were observed for other astrocytic states. The homeostatic state astH0 (enriched in synaptic-transmission and ion-homeostasis genes) was enriched among upregulated DEGs (adjusted p < 0.001) in non-carriers. Carriers showed enrichment of the astIM state (an intermediate transitional state between homeostatic and reactive astrocytes) among upregulated DEGs (adjusted p < 0.001). This state was not significantly enriched in non-carriers. Downregulated DEGs (adjusted p < 0.001) in carriers were enriched in the metabolic/chaperone-stress state astMet (enriched in heat-shock and energy-metabolism genes) and the intermediate state astIM.

**Fig. 6.**
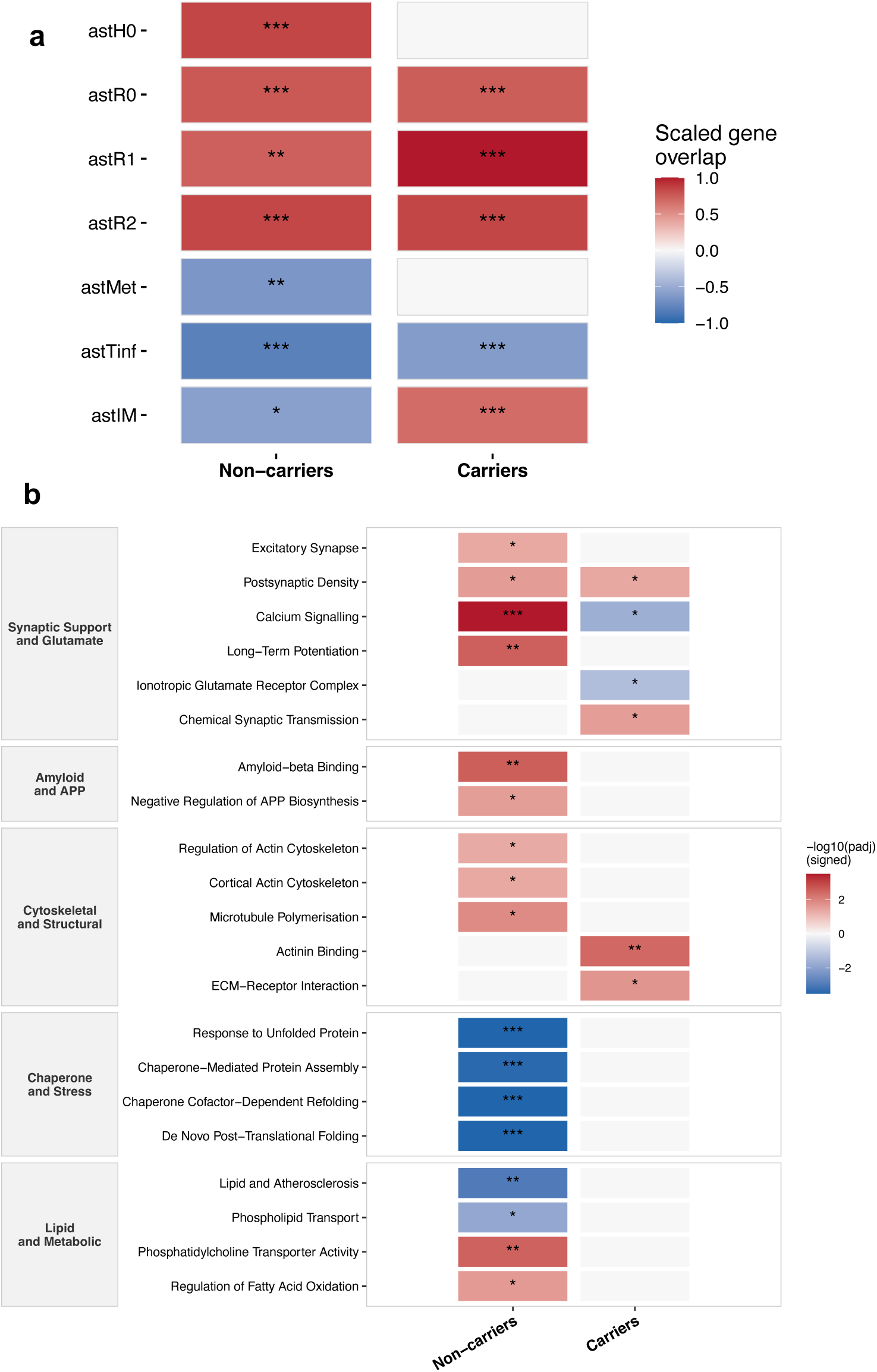
Differentially expressed genes with increasing amyloid-β load are associated with differences in astrocyte state and pathway enrichment for ABCA7 rs3752231 carriers and non-carriers. Enrichment analyses of significant astrocyte differentially expressed genes (DEGs; adjusted p ≤ 0.05, |log₂FC| ≥ 0.25) in ABCA7 rs3752231 carriers and non-carriers separately. Red tiles indicate upregulated enrichment; blue tiles indicate downregulated enrichment; white tiles indicate non-significant results. Asterisks denote significance thresholds: * adjusted p < 0.05, ** adjusted p < 0.01, *** adjusted p < 0.001. a Over-representation analysis (ClusterProfiler) of astrocyte DEGs for astrocyte state gene sets defined previously ^25^. Tile colour represents scaled gene overlap signed by direction of regulation, ranging from −1.0 (fully downregulated overlap) to +1.0 (fully upregulated overlap). All tested states are shown. b Pathway enrichment analysis of astrocyte DEGs based on Gene Ontology Biological Process, Cellular Component, Molecular Function, and KEGG 2021 Human pathway databases (EnrichR). Tile colour represents −log₁₀(FDR) signed by direction of enrichment.

Consistent with this, pathway enrichment analyses (DEGs; adjusted p ≤ 0.05, |log₂FC| ≥ 0.25) revealed largely genotype-specific patterns of enrichment (Fig. 6b). In non-carriers, astrocytes engaged a coordinated, multi-axis programme spanning synaptic support, amyloid handling and structural plasticity. Synaptic pathways, including excitatory synapse (*SRPX2, SYT11, SHISA6*), postsynaptic density (*GRIA1, GRIN2A, FXR1*), long-term potentiation (*CAMK2A, GRIA1, GRIN2A*) and calcium signalling (*CAMK2A, ITPR2, PRKCA*) were upregulated, as previously described for active astrocytic regulation of the tripartite synapse^30^. Amyloid-β binding (*CLU, APOE, LDLR*) and negative regulation of APP biosynthesis (*BACE2, ITM2B, ITM2C*) and cytoskeletal remodelling pathways (*GSN, ACTB, ARHGEF6*) also were enriched. the chaperone and unfolded protein response was strongly downregulated (Response to Unfolded Protein: *HSP90AA1, HSPA8, HSPH1*; padj < 0.001). This direction indicating proteostasis collapse in astrocytes with increasing Aβ burden is consistent with research reporting an overlapping gene set downregulated in astrocytes at high pathology states and interpret it as exhaustion of the chaperone response^25^. Fatty acid oxidation (*PPARA, PPARGC1A, ACADVL*) was selectively upregulated. We found a different pathway enrichment signature in carriers. Synaptic pathway enrichment appeared discordant, suggesting transcriptional dysregulation: postsynaptic density (*GRIA1, GRIN2C, GRIA3*) and modulation of chemical synaptic transmission (*GRIA1, GRM3, GRM8*) pathways were upregulated, but ionotropic glutamate receptor complex (*GRIA1, GRIN2C, GRIA3*) and calcium signalling (*CACNA1A, GRIN2C, PLCB1*) pathways were downregulated. In place of cytoskeletal plasticity, carrier astrocytes upregulated structural-matrix programmes (actinin binding: *CSRP1, SYNPO2, TTN*; ECM-receptor interaction: *TNC, FN1, ITGA2*), consistent with a shift from dynamic synaptic engagement towards matrix deposition. Amyloid-β binding, APP regulation, cytoskeletal plasticity, lipid handling and the chaperone response pathways were not significantly enriched. Together with the reduced glutamate transporter expression identified in the DEG analysis^31^, these findings suggest that carrier astrocytes do not show the range of potentially adaptive responses to Aβ pathology found in non-carriers.

### *ABCA7* genotypes are associated with differences in predicted microglia-astrocyte ligand-receptor interactions

Having identified genotype-specific transcriptional programmes in both microglia and astrocytes, we next explored differences in intercellular communication associated with these cell-intrinsic changes. We applied LIANA+ (ligand-receptor interaction analysis) as an exploratory framework to infer predicted microglia-astrocyte communication networks for each genotype. LIANA+ was applied with four aggregated scoring methods (Connectome, log2FC, NATMI, SingleCellSignalR) across all 54 donors (Fig. 1a), ranked by magnitude score separately within each genotype group (Fig. 7). Shared top-ranked interactions included those for microglial *C3*-*LRP1*/*NRP1*, *SPP1*-*CD44*, and *TGFB1* signalling to astrocytes, and for astrocyte *APOE*-*SORL1*, *TGFB2*-*TGFBR1*/*2*, and *HMGB1*-*TLR2* signalling to microglia. These transcriptional ligand-receptor pairs suggest shared intercellular communication features across genotypes (Fig. 7a). In non-carriers, the highest-ranked interaction was microglial *PDCD1LG2*-*RGMB*, a regulatory immune checkpoint signalling transcriptional pair associated with proteins underpinning the DAM-associate phagocytic responses to Aβ^32^ not expressed in carriers (Fig. 7b). Non-carriers also upregulated genes for of five astrocyte-derived ligands (*TF*, *EGF*, *HSPA8*, *VEGFB*, and *ARPC5*) that signal through the microglial β2-adrenergic receptor (*ADRB2*), expression of which in microglia modulates microglial inflammatory state and phagocytic responsiveness^33^. These receptor-ligand transcriptional pairs were not expressed in carriers.

**Fig. 7.**
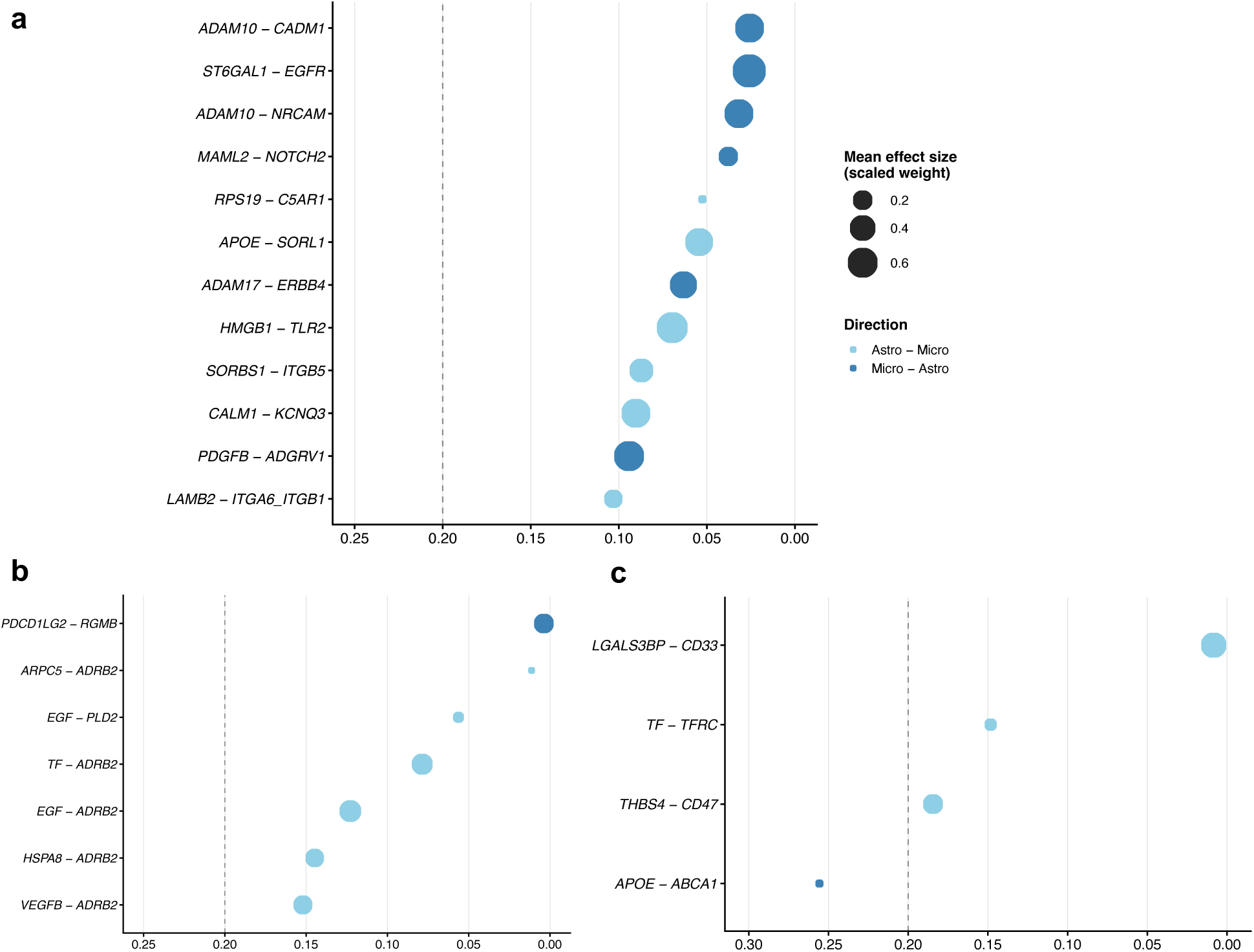
ABCA7 rs3752231 carriers show distinct patterns of transcriptional expression related to microglia-astrocyte ligand-receptor interactions compared to non-carriers. Ligand-receptor interaction analysis between microglia and astrocytes (LIANA+) with four aggregated scoring methods across 54 donors. Point position along the x-axis indicates magnitude rank (higher rank towards zero); point size reflects scaled interaction weight; light blue points indicate astrocyte→microglia signalling direction, dark blue points indicate microglia→astrocyte signalling direction. Dashed lines indicate magnitude rank = 0.20. Interactions were ranked by magnitude score within each genotype group. a Top-ranked ligand-receptor interactions shared between carriers and non-carriers within the top 50 ranked interactions of each group. b Top-ranked interactions unique to non-carriers. c Top-ranked interactions unique to carriers. Rankings are descriptive and should be considered hypothesis-generating.

Transcripts for carrier-specific interaction pairs included those related to phagocytosis and lipid metabolism (Fig. 7c). The highest-ranked astrocyte-to-microglia transcriptional interaction pair in carriers was for *LGALS3BP*-*CD33*. Galectin-3 binding protein signals through CD33, an inhibitory receptor that suppresses microglial phagocytosis that has been linked to AD risk^6,34^. Transcripts for astrocyte *THBS4*-*CD47* signalling to microglia, a ligand-receptor pair in which *CD47* acts as a “don’t eat me” signal that directly inhibits microglial phagocytosis via SIRPα also are upregulated^35^. Carriers also showed evidence for potentially adaptive upregulation of transcripts related to microglial *APOE*-*ABCA1* signalling to astrocytes, an interaction previously shown to regulate cholesterol efflux and lipid metabolism^36,37^.

## Discussion

Relative to homozygotes for the predominant *ABCA7* allele, heterozygotic carriers of the hypomorphic *ABCA7* rs3752231 variant showed greater total Aβ burden and greater diffuse plaque coverage in association with a lower proportion of compact plaques in late AD. This pattern is consistent with either impaired microglial-mediated plaque maturation or increased Aβ production. Using quantitative histopathology and single-nucleus transcriptomics on human *post-mortem* MTG, we provided transcriptomic evidence for the former by showing that the risk variant for AD is associated with suppressed canonical microglial activation and disrupted astrocyte homeostatic identity signatures. In addition to this evidence for impairment of microglial autonomous responses, we discovered that genes related to microglial-astrocyte signalling interactions expected to inhibit microglial phagocytic responses to Aβ also were upregulated^38^. We hypothesise that this functional pathology in microglia contributes to earliest AD pathogenesis in carriers.

A direct clinical implication of the work is for stratification of patients treated with anti-amyloid antibodies. Lecanemab and donanemab both rely on Fc-mediated microglial phagocytosis to clear opsonised Aβ. Our results provide evidence for an attenuated phagocytic response in rs3752231 carriers that predicts blunted efficacy of these antibodies in carriers. As about a third of the European AD population carries rs3752231, the impact of this could be significant. Testing for this should be prioritised.

Maturation of plaques in AD is an active process whereby microglia contribute to reshaping and compacting Aβ fibrils that limits Aβ neurotoxicity^16,20^. *TREM2* haplodeficiency in mice and humans impairs the microglial uptake and processing of Aβ central to this process, increasing relative proportions of diffuse plaques^39^. Reduced sequestration of neurotoxic Aβ fibrils by plaque compaction has been associated with greater neuronal dystrophy previously^17,18,40^. The similarities between the plaque endophenotypes and reduced microglial transcriptional responses to Aβ for *ABCA7* rs3752231 and *TREM2* R47H carriers^39,41^ thus provides evidence for related pathogenic mechanisms in human tissue in addition to any acceleration of Aβ production with ABCA7 hypomorphs through upregulation of the SREB2/BACE1 pathway ^42, 43,44,45^.

Microglial transcriptomic responses to Aβ pathology were markedly divergent between carriers and non-carriers of the *rs3752231* variant. Non-carriers upregulated expression of genes for canonical DAM markers (T*REM2*, *C1QA*, *C1QB*, *C1QC*, *AXL*, *APOE*) and genes for HLA antigen presentation as part of a previously well-characterised microglial phagocytic response that enhances the clearance of Aβ^22,46^. This signature was not found in carriers, which instead upregulated genes associated with a homeostatic state (*P2RY12*, *TGFB1*) while downregulating genes associated with cytokine (CRM2) and interferon response states (IRM)^22^. These findings conceptually map the functional consequences of this ABCA7 hypomorph onto impairment of the first, TREM2-independent stage of DAM activation, involving homeostatic checkpoint downregulation ^21^. *TREM2* hypomorphs exert risk by impairment of the second stage in this pathway. Part of *ABCA7* hypomorph risk for AD may be considered as arising through impairment of a major pathway shared with *TREM2* hypomorphs. We hypothesise that risks will be supra-additively expressed in carriers of both hypomorphic *ABCA7* and *TREM2* variants^21,47^.

Specific aspects of the molecular pathology for Aβ clearance by microglia were highlighted by our results. Differences in regulation of complement components (*C1QA*, *C1QB*, *C1QC*) in carriers and non-carriers is central to understanding why the rs3752231 variant enhances risks of AD. Increased C1q in early AD normally exerts protective effects by enhancing microglial clearance of Aβ oligomers, fibrillar plaques, and apoptotic neuronal debris^48^. Interaction of C1q with misfolded Aβ aggregates accelerates their nucleation into toxic ones and promotes clearance into fibrillar structures. This is central to plaque maturation in AD and a hallmark of the transition from early, diffuse deposits to mature, fibrillar, and neuritic plaques^49^. The contrasting upregulation of *C1Q* gene expression in non-carriers and its downregulation in carriers represents a direct molecular correlate of the divergence in phagocytic capacity between genotypes, a mechanism that would directly contribute to the observed increased accumulation of diffuse plaques^50^. A potential clinical implication is that therapeutic responses to anti-amyloid antibody therapeutics such as Lecanemab could be impaired in carriers of the common *ABCA7* rs3752231 variant^51^; stratification for this genetic variant could enhance the efficacy of this and related treatments.

Upregulation of expression of *SORL1*, *MGLL* and *MS4A6A* genes delineates an alternative microglial response to Aβ in rs3752231 carriers predicted to be associated with increased endosomal trafficking and lipid metabolic reprogramming rather than canonical immune activation. *SORL1* encodes a sorting receptor with an established role in microglial lysosomal homeostasis. Loss of SORLA in human iPSC-derived microglia impairs lysosomal enzyme trafficking and degradation of phagocytosed Aβ^52^; upregulation in carriers thus may reflect a compensatory response to impaired lysosomal processing of Aβ cargo in the context of reduced phagocytic activation*. SORL1* is also a negative regulator of *TREM2* surface expression and microglial lipid sensing^53^. *MGLL* encodes monoacylglycerol lipase, which hydrolyses the neuroprotective endocannabinoid 2-arachidonoylglycerol (2-AG); its upregulation reduces 2-AG availability and promotes pro-inflammatory lipid mediator production^47^. *MS4A6A* appears significantly downregulated in carriers. The gene is part of the *MS4A* gene cluster, several members of which constitute AD GWAS risk loci^54^. Reduced expression of the gene has been associated with exacerbated Aβ pathology and loss of microglial neuroprotective roles^55^. MS4A4A and MS4A6A also cooperate to negatively regulate TREM2 expression and microglial metabolism, phagocytosis, and lysosomal function, contributing to disease pathology^56,57^. Together, these findings frame a microglial response to increasing Aβ load in carriers distinct from the canonical DAM response in non-carriers^58^.

Astrocytes express similar *ABCA7* transcript levels to microglia^23^. Our work also identified genotype-specific differences in astrocyte responses to Aβ. Both *ABCA7* rs3752231 carriers and non-carriers upregulated canonical reactive transcriptional markers in astrocytes, but non-carriers also expressed homeostatic transcriptional pathways including those for synaptic support, glutamatergic signalling and long-term potentiation pathways, consistent with neuroprotective responses ^25,59,60^. It was notable that transcripts encoding glutamate transporters *SLC1A3* (EAAT1) and *SLC1A2* (EAAT2) were amongst those most downregulated in carriers. EAAT2 accounts for approximately 90% of glutamate uptake in the mature brain. Reduced EAAT2 activity impairs synaptic information transfer and leads to loss of excitatory-inhibitory homeostasis^27,61^. Loss of astrocyte-mediated synaptic support is an established contributor to neuronal vulnerability in AD^62^. In carriers, these functional effects would be expressed in the context of impaired clearance and sequestration of Aβ by microglia leading to increased levels of fibrillar forms of Aβ, which induce neuronal hyperexcitability^63^. This downstream consequence of impaired microglial function would be expected to be potentiated by the loss of astrocytic capacity to maintain synaptic homeostasis and contribute to increased neuronal vulnerability. Carrier-associated astrocytic pathology thus may potentiate these downstream neurodegenerative consequences, with astrocytes expressing *ABCA7* rs3752231 conferring increased vulnerability to excitotoxicity. Carriers of *ABCA7* hypomorphic variants - and particularly those who also are heterozygotic for *APOE4*, which has been implicated in Aβ-associated hyperexcitability in several earlier studies, may be a rational population to enrich in studies of neuroprotective treatments intended to reduce hyperexcitability^64^.

Both carriers and non-carriers increased expression of genes encoding ligand-receptor pairs for microglial *C3*-*LRP1*/*NRP1* and *SPP1*-*CD44* signalling to astrocytes, and astrocyte *APOE*-*SORL1* and *TGFB2*-*TGFBR1/2* signalling to microglia with increasing Aβ. However, our exploratory transcriptional ligand-receptor analysis identified evidence for genotype-specific patterns of microglia-astrocyte communication. For example, the increased expression of transcripts encoding five astrocyte-derived ligands for microglial *ADRB2* in non-carriers, whose activation modulates microglial inflammatory tone and phagocytic capacity^33,65^, was not found in carriers. This suggests loss of an astrocyte-mediated mechanism for sustaining microglial responsiveness with increasing Aβ in carriers. Carrier-specific ligand-receptor pairs were instead enriched for transcript pairs genes functionally associated with phagocytic suppression. The highest-ranked astrocyte-to-microglia interaction suggested for carriers, *LGALS3BP*-*CD33*, would functionally antagonise *TREM2*-mediated microglial activation through CD33 if expressed^34^. Evidence for *THBS4*-*CD47* signalling in carriers also predicts an additional phagocytic brake through the “don’t eat me” pathway^35,66^. These hypothesised interactions are consistent with independent experimental evidence that ABCA7 deficiency alters microglia–astrocyte communication^67^. Validation of specific interactions identified here by functional studies, would suggest further opportunities for therapeutic stratification, e.g., for carriers in assessing efficacy of adrenergic agonists^68^ or CD33 inhibitors.

A strength of our study lies in the joint characterisation of immunohistological and transcriptomic endophenotypes for the *ABCA7* rs3752231 allele in human tissue and with AD. However, there are limitations. First, the study is confined to donors from UK brain banks so cannot address the basis of differences in effects of *ABCA7* rs3752231 allele expression with different genetic backgrounds^69^. Second, the cross-sectional design precludes causal inference. Third, underrepresentation of mid-stage Braak III-IV donors may obscure the dynamic microglial transition window most relevant to DAM activation failure. We have, however, been able to identify genotype-dependent transcriptional responses to equivalent levels of Aβ burden, although the unequal group sizes (non-carriers n=19, carriers n=35) raise the possibility of differential sensitivities. A power analysis using a pre-specified minimum effect size (|log₂FC| ≥ 0.10) suggested that the current sample sizes should provide good levels of confidence in the primary results. To strengthen the robustness of pathway-level conclusions, state over-representation and pathway enrichment analyses (Figs. 4 and 6) also were performed at a stricter |log₂FC| ≥ 0.25 threshold than the |log₂FC| ≥ 0.10 used for gene-level analyses, prioritising larger effect sizes for gene-set composition. Fourth, the absence of spatial transcriptomic data means that the transcriptional impairments we observe cannot yet be localised to plaque microenvironments, which may be expected to differ widely^70^. Confirming that carrier-specific glial dysfunction is spatially coupled to Aβ would be an important focus for a follow-on study. Finally, the ligand-receptor analysis can only be considered as hypothesis generating; as noted above, the observations require functional validation in primary human microglia and astrocyte co-culture models with *ABCA7* variant background.

In summary, our work highlights that *ABCA7* rs3752231 carrier status is associated not merely with greater Aβ burden, but with failure of the coordinated glial response to Aβ and for plaque maturation. Diffuse plaque nucleation and growth, impaired microglial DAM activation and astrocyte homeostatic functions, downregulation of glutamate transporters, and convergent intercellular phagocytic suppression in the variant carriers all may contribute to the genetic risk conferred. Our findings establish *ABCA7* as a regulator of the integrated glial resilience to Aβ, operating through mechanism that partially converges with those of other risk factors involved in immune response and lipid metabolism, particularly *TREM2*. We hypothesised that *ABCA7* rs3752231 carriers, who are common amongst people with European ancestry, may show a reduced response to anti-amyloid treatments such as lecanemab or donanemab. We also could identify multiple complementary axes at which the carrier-specific functional impairments could be addressed therapeutically. Restoring intrinsic microglial activation in carriers might be achieved through TREM2 agonism, or through modulation of SORL1, which both restricts TREM2 receptor availability and is itself upregulated in carrier microglia. Removing extrinsic inhibitory signalling could be approached through CD33 inhibition. Finally, the carrier-specific reduction in astrocytic *SLC1A2* and *SLC1A3* supports investigation of EAAT2 potentiators to restore synaptic glutamate clearance and reduce excitotoxic neuronal vulnerability. Together, these observations position *ABCA7* rs3752231 not only as a risk variant but as a candidate stratification marker for several classes of potential future therapeutics.

## Methods

### Brain Tissue

Brain tissue acquisition, handling and processing were conducted under Human Tissue Authority licence number 12275, held by Imperial College Healthcare NHS Trust, in accordance with the Human Tissue Act 2004. Donor brains were obtained from six UK brain banks: the London Neurodegenerative Diseases Brain Bank (King’s College London), the Newcastle Brain Tissue Resource, the Queen Square Brain Bank (UCL), the Manchester Brain Bank, the Oxford Brain Bank, and the Parkinson’s UK Brain Bank (Imperial College London). Cases were selected based on neuropathological diagnosis of Alzheimer’s disease, spanning Braak stages 0–VI where possible, and the presence or absence of the *ABCA7* rs3752231 risk variant. No homozygous carriers of the variant were found in the cohort; carriers were heterozygous for the rs3752231 variant, non-carriers carry the common variant. Individuals with evidence of co-existing stroke, diabetes, Lewy body pathology, or TDP-43 pathology were excluded. In total, the cohort comprised mid temporal gyrus (MTG) tissue from 102 individuals, obtained as paired fresh frozen (FF) and formalin-fixed paraffin-embedded (FFPE) preparations from contralateral hemispheres, where available. 99 samples underwent 4G8 immunostaining of FFPE tissue for Aβ plaque characterisation, while 54 (Fig. 1a) provided FF tissue for glial enrichment and single-nucleus RNA sequencing (snRNA-seq).

### Genotyping

A subset of the samples were genotyped as follows. DNA was extracted from the FF tissue using the Qiagen DNAeasy Blood and Tissue kit. Samples were genotyped using the UK Biobank Axiom Array with a subset confirmed on the Illumina NeuroBooster Array (NBA). Standard quality control procedures were applied at both the individual sample and the SNP level using PLINK (v1.9)^71^ and R (4.2.2)^72^. Relevant variants at the *ABCA7* and *APOE* loci were extracted to determine genotype for each sample. Genotyping data for the remaining cases were obtained through the UK Brain Banks Network, and the Cambridge Study^73^ in which DNA was extracted from brain tissue prior to exome sequencing (Illumina Nextera 62 Mb capture) with variants called by FreeBayes^74^; copy number variant (CNV) analysis (Illumina HumanOmniExpress-12 BeadChip); C9orf72 repeat expansion detection; and *APOE* genotyping.

### Immunohistochemistry (IHC) and image acquisition

IHC was performed on FFPE tissue sections from the MTG (8 μm thickness). Standard immunohistochemical procedures were followed using the ImmPRESS HRP Horse Anti-Mouse IgG Polymer Detection Kit in conjunction with the ImmPACT DAB Substrate Kit, according to the manufacturer’s instructions (Vector Laboratories). Sections were baked at 60°C, dewaxed, and rehydrated. Endogenous peroxidase activity was quenched with 0.3% hydrogen peroxide. Antigen retrieval was performed by steaming in ethylenediaminetetraacetic acid (EDTA) buffer (pH 9.0). Sections were incubated overnight with a primary antibody against Aβ (clone 4G8, BioLegend, cat. no. 800711; 1:10,000). Signal detection was achieved using the HRP secondary antibody followed by 4-minute DAB exposure. Sections were counterstained with Mayer’s haematoxylin, dehydrated, cleared, and coverslip mounted. Dried sections were digitised using the Leica Aperio AT2 brightfield scanner (Leica Biosystems) at 20× magnification at the Imperial College Brain Bank Microscopy Facility. Representative images are shown in Fig. 1bi and 1bii.

### Plaque characterisation and quality control

Aβ-stained whole-tissue images of the MTG were analysed for plaque morphology assisted by the use of an in-house trained artificial intelligence model implemented within the BioDock^75^ platform. A total of 152 labelled 2×2 tiles were used to train a deep learning object detection model employing a hybrid convolutional and recurrent neural network (CNN-RNN) architecture to detect morphological and border-related features. Features derived from these components were fused and passed through a fully connected prediction head to assign plaque subtype labels. Non-maximum suppression was applied to remove redundant detections, and a confidence threshold of 0.6 was selected following systematic evaluation of multiple thresholds to optimise the balance between sensitivity and specificity on annotated training tiles. Outputs were exported in JSON format containing subtype and detection metadata for each detected object. JSON outputs were imported into QuPath^76^ alongside whole-slide images, enabling visualisation of plaques and annotations within their tissue context. A region of interest (ROI) encompassing the MTG was manually selected and quality-controlled for annotation accuracy, by a researcher blinded to donors’ metadata, including genotype and disease stage. Aβ plaques were classified into three morphological subtypes (Fig. 1c): diffuse plaques, characterised by loosely distributed Aβ (i); compact plaques, characterised by a concentrated, roughly circular deposit (ii); and dense-core plaques, characterised by a compacted Aβ core surrounded by a diffuse “halo” (iii). Following manual quality control, ROI-level metrics and individual plaque annotations were exported for analysis in R. To avoid capturing intracellular amyloid deposits, objects with a diameter of 10μm or less were excluded from downstream metrics (assuming circular shape). Mean plaque size was computed for all plaques and per subtype. Total plaque coverage and per-subtype coverage were calculated as the proportion of ROI area occupied by plaques of each morphological category, defined as: 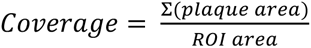. Total plaque coverage is used to illustrate Aβ burden in this study.

### Single nucleus isolation and selective glial enrichment for snRNAseq

Approximately 200 mg of FF MTG tissue was cryosectioned at 80 μm and collected on dry ice in BSA-coated DNA LoBind tubes. Nuclei isolation was optimised from a previously described protocol^77^. RNase-free conditions and 4°C temperature were maintained throughout. Briefly, tissue sections were mechanically homogenised using a glass dounce homogeniser in a homogenisation buffer containing 0.1% Triton X-100, RNase inhibitor (1 U/μL; Promega, N2511), protease inhibitor cocktail (Promega, G6521), and dithiothreitol (DTT). Myelin and cellular debris were removed by centrifugation through a sucrose-based density gradient (OptiPrep, Sigma-Aldrich). Nuclear integrity was confirmed by brightfield microscopy prior to sorting. Fluorescence-activated nuclei sorting (FANS) was performed on a BD FACS Melody instrument. Nuclei were stained with DAPI and antibodies targeting the pan-neuronal marker NeuN (Millipore, MAB377; 1:500) and the oligodendrocyte marker SOX10 (R&D Systems, AF2864; 1:250), followed by species-appropriate fluorescent secondary antibodies. The DAPI-positive, NeuN-negative, SOX10-negative population was collected for downstream snRNA-seq.

### Single nucleus capture, processing and sequencing

Following nuclei isolation and glial enrichment, snRNA-seq libraries were prepared using the Chromium Single Cell 3′ Gene Expression v3.1 Kit (10x Genomics) according to the manufacturer’s protocol. Nuclei were quantified using an acridine orange/propidium iodide stain on a Luna-FL Dual Fluorescence Cell Counter and loaded onto the Chromium Controller with a target recovery of 9,000 nuclei per sample. Individual nuclei were encapsulated into gel beads-in-emulsion (GEMs), where nuclear RNA was reverse-transcribed using barcoded oligonucleotides incorporating cell-specific barcodes and unique molecular identifiers (UMIs). cDNA was amplified (10 cycles), and sequencing libraries were generated following standard 10x Genomics workflows (14 cycles of final amplification).

Library quality was assessed using the Qubit dsDNA HS Assay Kit (ThermoFisher Scientific, Q32851) and the Bioanalyzer High Sensitivity DNA Kit (Agilent, 5067-4627). Libraries were sequenced at a depth of 50,000 reads per nucleus on the Illumina NovaSeq6000 platform at the Imperial College London Genomics Facility.

### Pre-processing and quality-control of snRNAseq data

FASTQ files were aligned to the pre-mRNA GRCh38 human reference genome using 10x Genomics Cell Ranger (v7.1.0). Ambient RNA contamination was removed from the raw feature-barcode matrices using CellBender (v0.03.0) (remove background) with default parameters. The CellBender output filtered feature-barcode matrices were used for downstream primary analyses using scFlow^78^, an in-house pipeline (v0.7.5). Quality control thresholds were applied as follows: library size 500–20,000 counts; number of detected features 400–8,000; mitochondrial gene fraction ≤10%. To prevent overrepresentation of any individual donor, samples contributing more than 3 standard deviations above the mean in cell count, mitochondrial read fraction, or total features by counts were excluded. Gene expressivity was defined as a minimum of 2 counts in at least 3 nuclei. Doublets were identified using DoubletFinder^79^ with a doublets-per-thousand-cells increment of 8, a pK value of 0.005, and embeddings generated from the first 10 principal components computed from the top 2,000 most highly variable genes (HVGs).

### Integration and clustering

Dataset integration across samples was performed using the LIGER package^80^(Linked Inference of Genomic Experimental Relationships), which factorises the gene expression matrix across samples into shared and sample-specific components. The following parameters were used: k = 35, λ = 5.0, convergence threshold = 0.0001, maximum iterations = 100, k-nearest neighbours k = 20, minimum cells = 2, quantiles = 50, random starts = 10, resolution = 1, number of genes = 3,000, centring = FALSE. Two-dimensional embeddings of the LIGER-integrated factors were computed using the Uniform Manifold Approximation and Projection (UMAP) algorithm^81^ with the following parameters: PCA dimensions = 30, n neighbours = 50, initialisation = spectral, metric = Euclidean, epochs = 200, learning rate = 1, minimum distance = 0.4, spread = 0.85, set operations mix ratio = 1, local connectivity = 1, repulsion strength = 1, negative sample rate = 5, fast SGD = FALSE. Unsupervised community detection was performed using the Leiden algorithm^82^ with a resolution of 0.0001 and k = 35. Cell-type annotation was performed using Expression Weighted Celltype Enrichment (EWCE) ^83^ within scFlow, against a reference dataset derived from the Allen Human Brain Atlas^7^. Automated annotations were subsequently reviewed by manual inspection of top cluster marker genes against canonical cell-type markers.

### Refinement of cell-type populations

To ensure analysis of transcriptionally pure cell populations, microglia and astrocytes clusters were independently extracted from the full integrated SingleCellExperiment^84^ object and converted to Seurat v5^85^ objects, retaining donor and sample metadata. For each cell type, raw counts were used as input; libraries were log-normalised, and the top 2,000 highly variable features were identified using the variance-stabilising transformation (vst) method. Data were scaled and principal component analysis (PCA) was performed on variable features. To mitigate donor-specific batch effects, Harmony^86^ was applied using sample identity as the integration variable, and UMAP embeddings were computed from the Harmony-corrected principal components for visualisation. Graph-based clustering was performed on the Harmony neighbourhood graph (dims 1–20) using the Louvain^87^ algorithm across multiple resolutions (0.4, 0.5, 0.6).

Cluster marker genes were identified using Wilcoxon rank-sum tests (FindAllMarkers; positive markers only; min.pct = 0.25; log2FC threshold = 0.25), and results were visualised using UMAP projections and marker heatmaps. An initial round of clustering was performed to identify and remove contaminating cell populations (including neurons, oligodendrocytes, endothelial cells, and pericytes), identified through EWCE-based reference mapping^83^ and confirmed by manual inspection of marker gene profiles. Following removal of contaminants, the cleaned cell population was re-normalised, and the full dimensionality reduction and clustering pipeline was repeated to yield the final cell-type-specific datasets used for downstream differential gene expression analyses.

### Pseudobulk differential gene expression analysis

To increase statistical power while preserving donor-level biological variability, single-nucleus RNA-seq data were aggregated into pseudocells using a random pseudobulk approach adapted from reported method^88^. Within each donor, nuclei were randomly aggregated into pseudocells of mean size 30 (minimum 10 nuclei per pseudocell). Prior to differential expression testing, genes were retained if their summed counts per million (CPM) across pseudocells exceeded 10% of the total pseudocell count and if they were detected in more than 5% of pseudocells, ensuring that only reliably expressed genes were included in downstream analyses.

Differential expression was performed using limma-trend within the limma/edgeR^89,90^ framework. A linear model was fitted per gene with total Aβ plaque coverage (continuous) as the primary variable of interest. All models were adjusted for the following covariates: pseudocell size (scaled), mitochondrial transcript percentage (log2-transformed), library size (scaled UMI count), age at death, sex, *post-mortem* interval (PMI), and *APOE* genotype group. Donor identity was incorporated as a random effect using the duplicateCorrelation approach. Empirical Bayes moderation was applied with trend=TRUE and robust=TRUE to account for the mean-variance relationship inherent to pseudobulk data. For stratified analyses, performed independently within *ABCA7* risk variant carriers and non-carriers, models were fitted separately within each stratum using identical covariate structures. P-values were adjusted for multiple testing using the Benjamini-Hochberg (BH) method^91^. Differential expression results were thresholded in two tiers. For gene-level analyses (DEG identification, fold-change reporting, volcano plots) genes with padj ≤ 0.05 and |log2 fold-change| ≥ 0.10 were considered differentially expressed, reflecting the pre-specified minimum effect size of biological interest, given the continuous Aβ coverage model in which fold change is expressed per unit of plaque coverage. For downstream gene-set analyses (state over-representation analysis (ORA) and pathway enrichment), a stricter |log₂FC| ≥ 0.25 threshold was applied to strengthen the robustness of enrichment conclusions to gene list composition.

To evaluate whether observed group differences reflected differential statistical power rather than biological signal, post-hoc power and minimum detectable sample sizes were estimated for key genes. For each gene, the effect size was derived from the moderated t-statistic returned by limma, and statistical power to detect the pre-specified minimum effect size of biological interest (|log₂FC| ≥ 0.10) was estimated using a parametric bootstrap (1,000 iterations), simulating replicate studies under the estimated effect and residual variance at the relevant sample size (non-carriers n = 19, carriers n = 35). Minimum sample size was determined by increasing n until bootstrap-estimated power reached 80%.

### Over-representation analysis against microglial state gene sets

To determine whether differentially expressed microglial genes were enriched within known microglial transcriptional states, ORA was performed using the enricher function from ClusterProfiler^92^. Significant DEGs (padj ≤ 0.05 and |log₂FC| ≥ 0.25) were stratified by direction of effect (upregulated and downregulated) and tested separately against a curated set of microglial state gene sets defined previously^22^, comprising homeostatic (HM), disease-associated (DAM), antigen-presenting (DAM-HLA), cytokine-response (CRM), interferon-response (IRM), and perivascular (PVM) microglial states. The full set of genes detected across pseudocells was used as the statistical background universe. Enrichment significance was assessed using a hypergeometric test and p-values were adjusted using the BH method. An additional q-value cutoff of 0.2 was applied as implemented in ClusterProfiler, which estimates q-values using the Storey method. Results were visualised as a signed gene overlap heatmap, where tile colour reflects scaled gene overlap signed by direction of enrichment (upregulated = positive, downregulated = negative), and significance is indicated by asterisks (*, padj < 0.05; **, padj < 0.01; ***, padj < 0.001). An equivalent analysis was performed for astrocyte DEGs, tested against curated astrocyte transcriptional state gene sets^25^, comprising reactive (astR0, astR1, astR2), homeostatic (astH0), metabolic (astMet), inflammatory (asTInf), and immune-modulatory (astIM) states.

### Pathway enrichment analysis

Functional pathway enrichment analysis was performed on significant microglial and astrocyte DEGs (padj ≤ 0.05, |log₂FC| ≥ 0.25) using the EnrichR API^93^ via a custom wrapper function implementing the enrichR R package^94^. Separate analyses were performed for *ABCA7* risk variant carriers and non-carriers. The following databases were queried: GO Biological Process 2023, GO Cellular Component 2023, GO Molecular Function 2023^95^, and KEGG 2021 Human^96^. Gene sets were filtered to include only those with between 5 and 300 members and with at least 2 overlapping genes. P-values were adjusted for multiple testing using the BH method within each database, and gene sets with FDR ≤ 0.05 were retained. Pathway directionality was inferred by computing the mean log2 fold-change of overlapping genes, with positive and negative mean values assigned as upregulated and downregulated respectively. Results were visualised as a signed -log10(FDR) heatmap, with pathways grouped into biological themes: phagocytosis and antigen presentation, cytokine and inflammatory signalling, JAK-STAT and interferon signalling, cytoskeletal organisation and migration, and amyloid, lipid and stress responses for microglia and synaptic support & glutamate, amyloid and APP, cytoskeletal & structural, chaperone & stress and lipid & metabolic for astrocytes.

### Ligand-receptor interaction analysis

Ligand-receptor interaction analysis between microglia and astrocytes was performed using LIANA+^97^ with four scoring methods, Connectome, log2FC, NATMI, and SingleCellSignalR, applied independently to each donor using the rank_aggregate_by_sample function. Per-method scores were aggregated into unified magnitude and specificity ranks using robust rank aggregation (RRA) with the consensus ligand-receptor resource. Micro®Astro and Astro®Micro interactions were extracted separately for carriers and non-carriers. Interactions were assessed qualitatively rather than through formal differential testing, given the exploratory nature of this analysis, comparing the top-ranked Micro↔Astro interactions between carriers and non-carriers to identify biologically relevant differences in glial communication patterns.

## Acknowledgements

The authors thank Megan Winterbotham for outstanding laboratory management and support throughout this study. We thank the donors and their families for the use of human brain tissue in this study. Tissue samples were provided by the following brain banks, to whom we are grateful: the London Neurodegenerative Diseases Brain Bank at King’s College London, funded by the UK Medical Research Council and the Brains for Dementia Research programme, jointly funded by Alzheimer’s Research UK and the Alzheimer’s Society; the Newcastle Brain Tissue Resource, funded in part by a grant from the UK Medical Research Council (G0400074), NIHR Newcastle Biomedical Research Centre and Unit awarded to the Newcastle upon Tyne NHS Foundation Trust and Newcastle University, and the Brains for Dementia Research programme; the Queen’s Square Brain Bank, UCL; The Manchester Brain Bank, part of the Brains for Dementia Research programme, jointly funded by Alzheimer’s Research UK and Alzheimer’s Society; the Oxford Brain Bank, supported by the Medical Research Council, the NIHR Oxford Biomedical Research Centre, and the Brains for Dementia Research programme; and the Parkinson’s UK Brain Bank at Imperial, funded by Parkinson’s UK, a charity registered in England and Wales (258197) and in Scotland (SC037554). The Imperial BRC Genomics Facility was supported by the National Institute for Health Research Biomedical Research Centre.

## Author Contributions

Conceptualisation: P.M. Matthews, M. Papageorgopoulou. Data curation: M. Papageorgopoulou. Formal analysis: M. Papageorgopoulou. Funding acquisition: P.M. Matthews. Investigation: M. Papageorgopoulou, E. Adair, M. Wülfing. Methodology: M. Papageorgopoulou, N. Fancy, B. Avot, S. Boulger. Project administration: M. Papageorgopoulou, P.M. Matthews. Resources: P.M. Matthews. Supervision: P.M. Matthews. Visualisation: M. Papageorgopoulou. Writing, original draft: M. Papageorgopoulou. Writing, review and editing: M. Papageorgopoulou, P.M. Matthews.

